# Rho Kinase regulates neutrophil NET formation that is involved in UVB-induced skin inflammation

**DOI:** 10.1101/2021.05.16.444366

**Authors:** Minghui Li, Xing Lyu, James Liao, Victoria P. Werth, Ming-Lin Liu

## Abstract

**Objective:** Ultraviolet B (UVB) is an important trigger of skin inflammation and lupus with leukocyte recruitment to inflamed skin. We recently reported the involvement of neutrophil NETosis in UVB-induced skin inflammation, and that NETotic nuclear envelope rupture is driven by PKCα-mediated nuclear lamin B disassembly. To address the role of Actin cytoskeleton in NETosis, we investigated the effects of Rho kinase (ROCK) and its downstream actomyosin cytoskeletal networks on PKCα nuclear translocation and NET formation, as well as their involvement in UVB-induced skin inflammation.

**Methods:** We studied the dynamic changes of ROCK and actomyosin cytoskeletal networks during NETosis induction and their involvement in PKCα nuclear translocation. Using mice with hematopoietic-specific ROCK1 deficiency, we investigated the effects of ROCK1 deficiency on NETosis, and its involvement in UVB-induced skin inflammation.

**Results:** Our time course studies demonstrated the dynamic changes of actin polymerization and ROCK activation, support the role of actin cytoskeleton in nuclear translocation of cytosolic PKCα in early stage of NETosis induction. Inhibition of actin polymerization or key molecules of the ROCK/MLCK/myosin pathway decreased PKCα nuclear translocation and NET formation. Genetic deficiency of ROCK1, inhibited NETosis *ex vivo* and *in vivo*, decreased extracellular display of NET-associated IL-17A, TNFα, IFNγ, and IFNα in inflamed skin, which were correlated with the ameliorated skin inflammation in UVB-irradiated mice with hematopoietic-specific ROCK1 deficiency.

**Conclusions:** ROCK regulated NETosis through modulation of PKCα nuclear translocation via actomyosin cytoskeletal networks in neutrophils. ROCK1 deficiency ameliorated UVB-induced skin inflammation by attenuation of NETosis and NET-associated cytokines.

## INTRODUCTION

Ultraviolet B (UVB) is an important trigger of cutaneous and systemic lupus erythematosus (CLE, SLE) [1–4]. UVB overexposure causes photodamage of skin with recruitment of inflammatory cells, including neutrophils [5]. A recent study has indicated that neutrophils are the critical early infiltrating immune cells within cutaneous lupus lesion in mice [6]. Several studies with transcriptional profiling analysis of lupus patient samples also confirmed the importance of neutrophils [7], and a gradual enrichment of neutrophil transcripts during lupus disease progression [8]. Therefore, it is important to understand the role of neutrophils in UVB-induced skin inflammation. NETosis is a unique form of neutrophil cell death with release neutrophil extracellular traps (NETs) [9, 10], which are involved in lupus pathogenesis [11, 12]. However, the involvement of NETosis in UVB-induced skin inflammation and the relevant cellular mechanisms are not well understood.

Nuclear chromatin forms the backbone of NETs, and thus nuclear envelope rupture is a required step for breaking the first physical barrier, nuclear chromatin release and extracellular NET formation [9, 10, 12]. We recently found that NETotic nuclear envelope rupture is driven by protein kinase C alpha (PKCα)-mediated phosphorylation and disassembly of nuclear lamin B [9] which is crucial to nuclear envelope integrity by forming highly organized meshworks surrounding the nuclear chromatin [13]. Furthermore, cyclin-dependent kinases (CDK)4/6 also control NETosis through regulation of lamin A/C phosphorylation [14]. Therefore, PKCα and CDK4/6 serve as NETotic lamin kinases [9, 10, 14] for regulation of nuclear lamina disassembly and NETotic nuclear envelope rupture.

PKCα exists diffusely in the cytosol of the unstimulated cells, and the kinase can be translocated to the nucleus during activation [15]. Schmalz et al indicated that nuclear translocation of cytosolic PKCα requires intact cytoskeleton [16]. As an upstream regulator of cytoskeleton, Rho kinase (ROCK) inhibition blocks nuclear translocation and activation of PKC [17]. We have shown that PKCα nuclear translocation is required for nuclear lamina disassembly and NET formation [9]. Furthermore, functional cytoskeleton is required for NETosis induction [18–21] in the early stage, while actin cytoskeleton disassembly in the late stage is involved in the plasma membrane rupture during NETosis [10, 22].

It has long been known that Rho [23] and its effector kinase ROCK [24] regulate actin cytoskeleton organization. In the current study, we sought to investigate the dynamic changes of actin cytoskeleton and its upstream regulator ROCK, as well as its involvement in PKCα nuclear translocation during NETosis induction. Furthermore, F-actin and myosin motors form actin-myosin (actomyosin) cytoskeletal networks that are involved in nucleocytoplasmic shuttling [25, 26]. The actomyosin cytoskeletal networks can be regulated by activation of myosin II and myosin regulatory light chain kinase (MLCK) through the ROCK/MLCK/myosin pathway [27]. Therefore, we also investigated the involvement of ROCK/MLCK/myosin pathway in PKCα nuclear translocation and NET formation *in vitro*. Importantly, we investigated the effects of ROCK1 deficiency on neutrophil NET formation *in vivo* and UVB-induced skin inflammation using mice with hematopoietic-specific ROCK1 deficiency.

## METHODS

### Mice

ROCK1 deficient mice ROCK1^+/-^ on a C57BL/6J background were generated by Liao’s laboratory as described before [28]. Since homozygous (ROCK1^-/-^) mice always die perinatally, heterozygous mice were used in this study. C57BL/6J wildtype (WT) and CD45.1 (B6.SJL- *Ptprc^a^Pepc^b^*/BoyJ) mice were purchased from Jackson Laboratory. All mice and their corresponding littermate controls were housed in a pathogen-free environment, and given food and water *ad libitum*. All animal experiments and animal protocols were approved by the Animal Care and Use Committee of University of Pennsylvania.

### Generation of hematopoietic deficient mice by bone marrow transplantation (BMT)

To generate mice with hematopoietic-specific ROCK1 deficiency, or their corresponding background control mice, BMT was conducted according to our established BMT techniques by following the approved animal protocol, as described with modification [29]. In brief, the BM hematopoietic stem cells (HSCs, >5x10^6^ cells) were harvested from the femur and tibia of donor ROCK1^+/-^ mice or their littermate WT (all 6-8-week-old, with CD45.2 congenic background). The donor HSCs (>5x10^6^ cells) from either ROCK1^+/-^ or WT mice were transplanted into sex-matched CD45.1 recipient mice (all 6-8-week-old) which were pre-irradiated sub-lethally with two equal doses of 550 cGy (*Total Body Irradiation* 1100 cGy) at intervals of 18 hours [30, 31] with a *Cs*-*137 Irradiator* [29]. Starting one-week prior to BMT, mice were given medicated acidified water with antibiotics (sulfamethoxazole and trimethoprim) until 6 weeks post BMT. Mice were given soften food and kept in an animal biosafety level 2 room immediately after irradiation and BMT for up to 6 weeks until UVB exposure. Peripheral blood chimerism was analyzed by flow cytometry analysis of the reconstitution rate of CD45.2 cells in the peripheral blood of CD45.1 recipient mice at 4 weeks post BMT as a functional assay of HSC fitness. Only successfully transplanted mice (over 95% reconstitution) were used in later animal experiments.

### UVB Exposure

The dorsal skins of CD45.1 mice with hematopoietic-specific ROCK1 deficiency (BMT- ROCK1^+/-^) or their corresponding control mice (BMT-WT) were irradiated or not (sham groups) by UVB (150 mJ/cm^2^/day) for 5 consecutive days under anesthetization by peritoneal injected Ketamine/Xylazine [9]. Each group included 6-8 mice with half males and half females. Animals were then sacrificed 24 hours after the last exposure. Whole dorsal skin samples, including epidermis, dermis, and subcutaneous fat were collected. Tissue sample sectioning was performed by the Penn Skin Biology and Diseases Resource-based Center. Skin sections from UVB-irradiated and sham mice were stained by hematoxylin and eosin (H&E). Infiltration of inflammatory cells was counted for the nucleated cells in the epidermis, dermis, or fat [32, 33].

### Cell Culture and Treatment

Primary human neutrophils from the peripheral blood or mouse bone marrow (BM) neutrophils from the *femur* and *tibia* BM of WT vs ROCK1^+/-^ [28] were isolated as described in our publication [9]. Mouse BM mPMNs were isolated with a neutrophil isolation kit (Miltenyl). Human HL-60 cell line were differentiated into polymorphic nuclear granulocytes (dPMNs) in RPMI-1640 with 10% FBS and 1.2% DMSO for 5-6 days as described in our published study [9]. A time course study of treatment of dPMNs by 50 nM PMA was conducted for 0, 15, 30, 60, 120, and 180 min, and then cells were either fixed with 2% paraformaldehyde, or lysed with cell lysis buffer, for different experiments. In separate experiments, the above primary human or mouse neutrophils or dPMNs were treated with, either PMA at 50-100 nM, or physiological NETosis- inducer platelet-activating factor (PAF) [9] at 5-10 µM, for 1h or 3h without or with 30 min pretreatment with inhibitors of F-actin polymerization (Cytochalasin D), myosin II (Blebbistatin), myosin light chain (MLC) kinase (ML7), or ROCK (Y-27632, HA-1077, AS1892802), following by detection of NET formation or actin polymerization. In the above time course studies or experiments using different inhibitors, the confocal microscopic imaging analyses, of the 0 time point, or the control conditions. started after 5 min of PMA stimulation to allow the suspension dPMNs to adhere to petri dish in order to compare them with the adherent dPMNs at either later time points of the time course, or other conditions without or with inhibitors.

Detection of ROCK activity was performed with a kit from Millipore. Briefly, neutrophil lysates from different time points were added to a plate coated with recombinant Myosin phosphatase target subunit 1 (MYPT1) that contains a Thr696 residue (MYPT1^Thr696^) which can be phosphorylated by ROCK1/2. A detection antibody that specifically detects MYPT1^Thr696^ was then applied, following by an HRP-conjugated secondary detection antibody, and chromogenic substrate. After adding stop solution, the absorbance signal detected at 450 nm reflected the relative amount of ROCK activity in the samples.

### Assessment of NET Formation

All of the above primary human or mouse neutrophils or dPMNs were treated with either PMA (50 or 100 nM) or PAF (5 or 10 µM) for 3h. NET formation assessment and quantification was analyzed by either fluorometric NET quantification by SYTOX Green staining intensity, and/or immunofluorescent imaging quantification analysis by staining with impermeable DNA dye SYTOX Green for detection of NET formation, and the cell-permeable DNA dye SYTO Red to count the total number of neutrophils, as described before [9].

### Flow cytometry analysis

For detection of polymerized filamentous actin (F-actin), the 2% paraformaldehyde (PFA) fixed neutrophils from different time points (time course) after PMA stimulation, or PMA-stimulated dPMN cells or mouse primary BM neutrophils without or with pretreatment by ROCK inhibitors were stained by phalloidin-RFP or phalloidin-FITC, and analyzed with flow cytometry according to published methods [34]. To detect the indirect effects of UVB through PAF [35] on cytokine induction, the BM neutrophils from ROCK1^+/-^ vs WT were treated overnight without or with PAF, without or with isolated NETs (for IFNα induction). Then the treated cells were fixed with 2% PFA and permeabilized with 0.1% Triton X-100, and stained by FITC- or Alexa488- conjugated primary antibodies against IL-17A, TNFα, IFNγ, or IFNα (Biolegend or Bioss). Actin polymerization was detected by RFP- or FITC-conjugated phalloidin. Expression of CXCR2 in BM neutrophils was detected by FITC-labeled anti-CXCR2. Expression of cytokines or CXCR2 and actin polymerization in mouse BM neutrophils were detected based on gated cells that were positive for neutrophil marker ly6G (PE-labeled). Furthermore, to detect the reconstitution rate of donor-derived cells in the peripheral blood of BMT mice, blood samples were harvested from the saphenous vein of all mice 4 weeks after the BMT procedure. Erythrocytes were then removed by hypotonic lysis (0.2% NaCl2), and stained with PE-labeled anti-mouse CD45.1 and FITC-labeled anti-mouse CD45.2, following by flow cytometry analysis as described before [36].

### Transmigration Assays

The pore transmigration ability of primary neutrophils from WT vs ROCK1^+/-^ mice was examined as described before [9] by using Boyden chamber, 24-well trans-well plates with 3 µm pore inserts, under stimulation with CXCL-1 (100 ng/ml). The pore transmigration ability was expressed as relative transmigration compared to that in control neutrophils of the mice without treatment.

### Fluorescent Immunocytochemistry and Immunohistochemistry analysis

For immuno-cytochemistry analysis, dPMNs or mouse BM neutrophils were treated without or with different inhibitors, then treated without or with PMA or PAF for 1-3h for study of PKCα nuclear translocation. Then the treated cells were stained by primary anti-total PKCα or anti-lamin B Abs, followed by secondary Abs with FITC or PE conjugation [9]. For the *ex vivo* NETosis induction with exhibition of NET-associated cytokines, the BM neutrophils from ROCK1^+/-^ vs WT were treated for 20h without or with PAF, without or with isolated NETs (for IFNα induction). Then the treated cells were fixed with 2% PFA and permeabilized with 0.1% Triton X-100, and stained by primary antibodies against IL-17A, TNFα, IFNγ, or IFNα with FITC- or Alexa488-conjugation (Biolegend or Bioss). The neutrophil marker Alexa-647-labeled anti- ly6G was used. DNA was stained with DAPI.

The paraffin-embedded skin tissue sections were stained as described before [9]. The neutrophil markers Alexa-647-labeled anti-ly6G were used for detection of neutrophils and NETosis in combination with DNA staining with DAPI as described [9]. FITC- or Alexa488- conjugated primary antibodies against IL-17A, TNFα, IFNγ, or IFNα were used to stain cytokines (Biolegend or Bioss). Slides were mounted with Gold Antifade Mountant (Invitrogen). Confocal fluorescent images were analyzed with an Olympus Fluoview 1000 confocal microscope.

### Statistical analyses

GraphPad Prism 6 was used to perform statistical analysis. Data are shown as means ± standard deviations (SD). The sample sets were first analyzed for normality. Comparisons amongst three or more groups were performed using ANOVA followed by Student-Newman-Keuls analysis for sample sets with normal distribution, or using Kruskall-Wallis analysis followed by Dunńs test for sample sets with a non-normal distribution. Comparisons between two groups were conducted with student’s t-test. Pearson correlation analysis was used when applicable. Statistical significance was considered at a level of *P*-value < 0.05.

## RESULTS

### Disruption of actin cytoskeleton or actomyosin cytoskeletal networks attenuated neutrophil NET formation

Nuclear envelope rupture is regulated by PKCα-mediated lamin B disassembly [9, 10], and nuclear translocation of PKCα requires intact cytoskeleton [16, 21, 37]. In addition, actomyosin networks have been shown to be important in nucleocytoplasmic shuttling [25, 26]. In the current study, we found that disruption of actin cytoskeleton filaments by inhibition of actin polymerization with cytochalasin D, or disruption of actomyosin cytoskeletal networks by inhibition of myosin II or its upstream MLCK with Blebbistatin or ML7 correspondingly, significantly decreased PMA- or PAF-induced NETosis in human or mouse neutrophils (Figure 1A-C). Importantly, disruption of actin cytoskeleton by cytochalasin D, or inhibition of myosin II or MLCK by Blebbistatin or ML7, increased cytosolic retention of PKCα and attenuated nuclear translocation of PKCα, observed at 60 min after PMA stimulation (Fig 1D-F). Interestingly, the nuclei of these cells still remained as multiple lobes similar as those in control cells (Fig 1D-F). In contrast, we saw the swelling nuclei with disintegrated nuclear envelope in cells treated with PMA alone (Fig 1D). These results suggested that not only actin cytoskeleton, but also the actomyosin cytoskeletal networks, are important for nuclear translocation of PKCα and nuclear envelope rupture (Fig 1G), which is required for nuclear chromatin extrusion and NET formation [9, 10]. Along this line, other studies also reported the involvement of actin cytoskeleton [38] or myosin II [39] in nuclear translocation of cytosolic molecules, and in actomyosin-dependent nucleocytoplasmic shuttling [25, 26].

**Figure 1.**
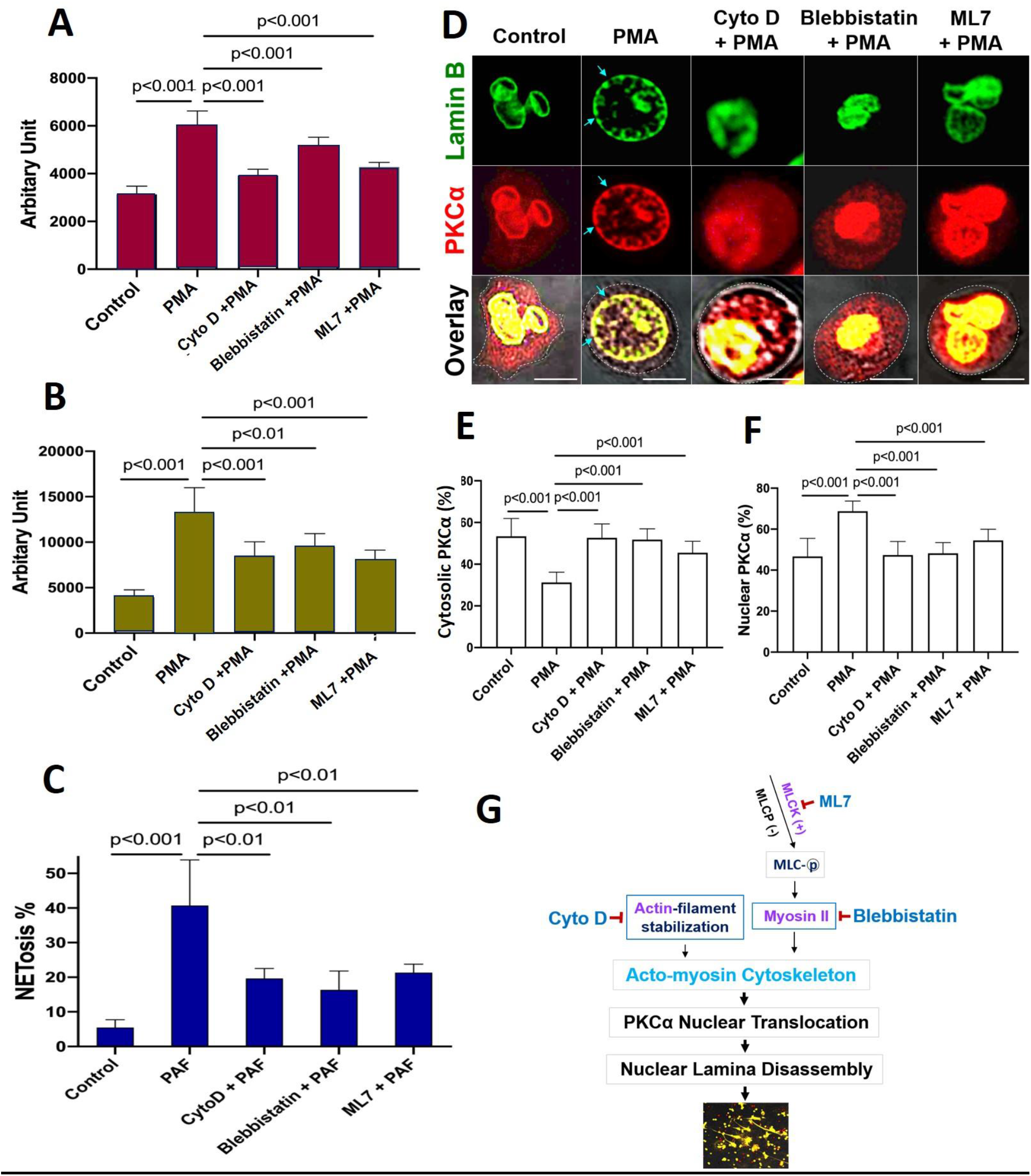
Disruption of actin cytoskeleton or actomyosin cytoskeletal networks attenuated NETosis in neutrophils. **A**,**B**. dPMNs (**A**) or primary human neutrophils from different healthy donors (**B**) were exposed to 50 nM PMA for 3h without or with 30 min pretreatment by 10 μM Cytochalasin D (Cyto D), 50 μM Blebbistatin, or 1 μM ML7, then detection of NETosis with Sytox Green staining by microplate reader. **C**. Primary bone marrow (BM) mouse neutrophils from different C57BL/6 wild type (WT) mice were treated with 10 μM PAF (a physiological stimulus) for 3h, without or with 30 min pretreatment by 10 μM Cyto D, 50 μM Blebbistatin, or 1 μM ML7, and stained by cell permeable SYTO Red and cell impermeable Sytox Green, then detected by fluorescent microscopy (**C**). Images were taken by confocal microscopy, followed by automated quantification of NETs in 5-6 non-overlapping areas per well using ImageJ for calculation of % cells with NET formation. **D-F**. Representative images and summary analysis for the effects of Cyto D, Blebbistatin, and ML7 on nuclear translocation of PKCα and nuclear lamina (lamin B) disintegration in dPMNs exposed to 50 nM PMA for 1h without or with pretreatment by above inhibitors. All cells were stained for **lamin B** and **PKCα** with corresponding antibodies. Scale bars, 10 μm. **G.** Schematic illustration of the involvement of actin cytoskeleton and actomyosin cytoskeletal networks, and their corresponding inhibitors in NETosis. All results in panels A-C represent 5-6 biological replicates. Panels E-F are summary analyses of at least 10 cells for each conditions from three independent experiments. For sample sets with normal distribution, ANOVA analysis with post hoc Student-Newman Keuls test were conducted. Panels **A-C,E,F** display means±SD.

### ROCK regulated PKCα nuclear translocation by modulation of actin cytoskeleton polymerization in neutrophils

To understand the potential role of actin cytoskeleton and its upstream regulator ROCK in PKCα nuclear translocation, we investigated the time course of PKCα nuclear translocation, the dynamic changes of actin polymerization, and Rho kinase activation in PMA-stimulated cells from 0 to 3h. As we have shown in our recent publication [9], the diffusely distributed cytosolic PKCα in PMA-stimulated cells gradually translocated to the nucleus within 60 minutes of PMA stimulation (Fig 2A-C), as compared to those at initial time point. Alone with the nuclear accumulation of lamin kinase PKCα, PKCα-mediated nuclear lamin B disassembly resulted in nuclear envelope disintegration at 60 min (Fig 2A), followed by nuclear DNA extracellular extrusion from small rupture sites at 120 min (Fig 2A), and the extracellular release of large amounts of nuclear DNA from the enlarged rupture of the nuclear envelope at 180 min (Fig 2A) after PMA stimulation, resulting in extracellular NET formation with the collapse of the nuclear and plasma membranes [9, 10].

**Figure 2.**
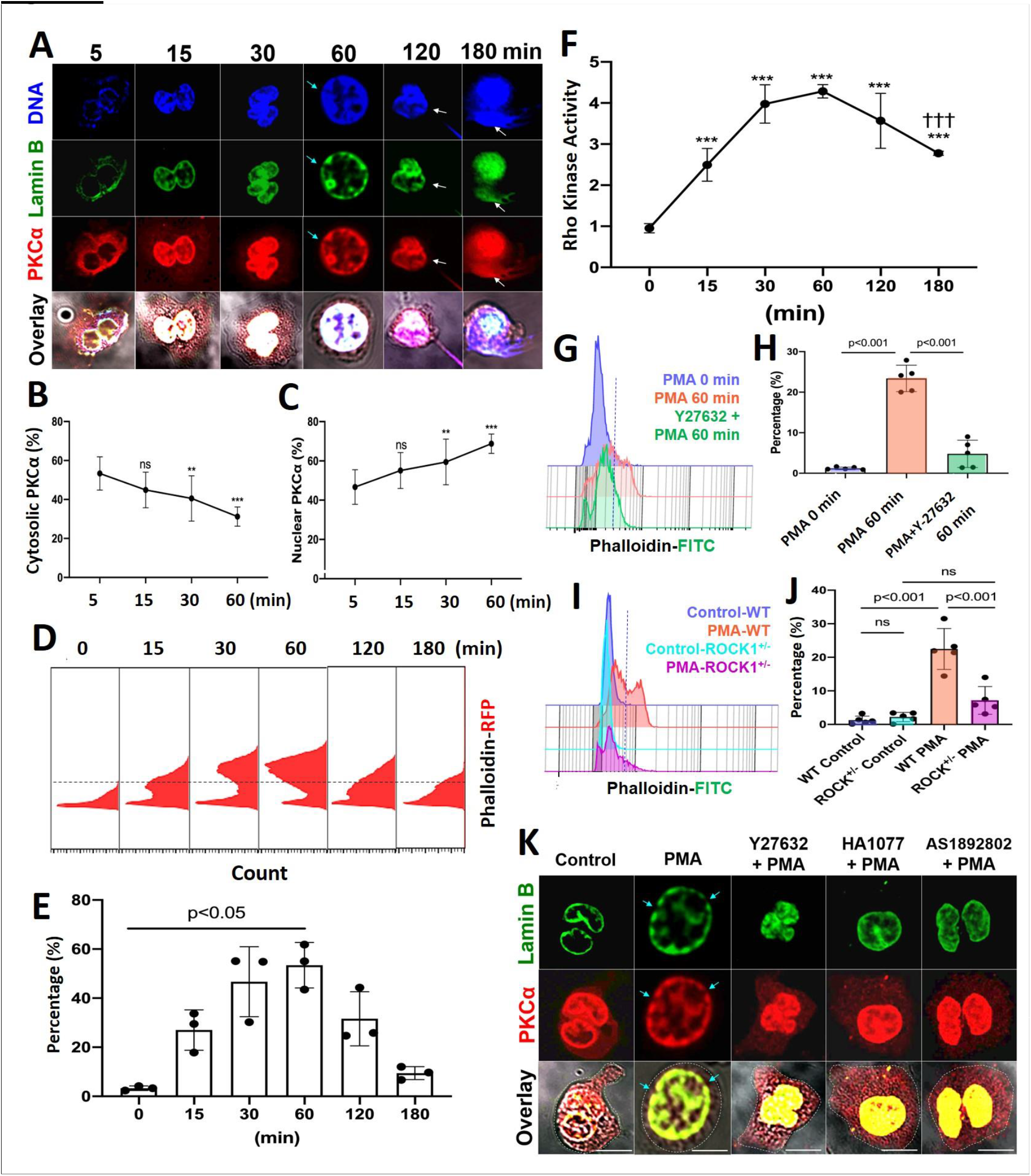
PKCα nuclear translocation and actin cytoskeleton assembly were regulated by Rho kinase 1 (ROCK1). **A.** Representative images for the time course study of PKCα nuclear translocation, subsequent lamin B disassembly and nuclear lamina disintegration, and NET release in dPMNs exposed to 50 nM PMA for 0-180 min, then stained simultaneously for **DNA**, **lamin B** (probed by primary anti-lamin B, and FITC-labeled secondary antibody), and **PKCα** (primary anti- PKCα, and PE-labeled secondary antibody) under confocal fluorescent microscopy. Scale bars, 10 μm. **B-C**. Summary analyses of panel A for cellular distribution of both cytosolic (B) and nuclear (C) PKCα in dPMNs treated with 50 nM PMA for 0-60 min. For 0 min at panels A-C, analysis started after 5 min of PMA stimulation to allow the suspension dPMNs to adhere in order to compare with the adherent dPMNs at later time points for microscopic analysis. **D-E**. Representative flow cytometry histograms and summary analyses for the time course of F-actin polymerization in dPMNs exposed to 50 nM PMA for 0-180 min, the pemeabilized cells were probed with phalloidin-RFP. **F**. Time course of ROCK activation in dPMNs exposed to 50 nM PMA for 0-180 min, detected by ROCK activity ELISA. **G-H**. Representative and summary analyses of F-actin polymerization in WT primary mouse neutrophils stimulated by 50 nM PMA for 0 or 60 min, and PMA treatment at 60 min without or with pretreatment by 50 μM ROCK inhibitors Y27632 for 30 min. **I-J**. Representative and summary analyses of F-actin polymerization in mouse neutrophils from WT or ROCK^+/-^ mice stimulated without or with 50 nM PMA for 60 min. Cells in panels G-J were detected by flow cytometry by staining with phalloidin-FITC and anti-ly6G-PE. **K**. Representative images for the effect of ROCK inhibition on PKCα nuclear translocation and nuclear envelope disintegration in dPMNs exposed to PMA for 60 min without or with pretreatment by ROCK inhibitors, control cells same as 0 min in panels A-C. All cells were stained for **lamin B** and **PKCα** with their corresponding Abs. Scale bar, 10 μm. Light blue arrows in panels A,K indicate the disintegration/rupture of nuclear lamina at 60 min after PMA treatment, while white arrows in panel A indicate the rupture site of the nuclear envelope where the decondensed chromatin released for NET formation at later time points. Panels B-C were summary analyses of at least 10 cells at each time point (5-60 min) from three independent experiments. The results of summary analyses represent 3 (E), 4 (F), or 5 (H,J) biological replicates. For non-normal distributed data (E), Kruskall-Wallis analysis and Dunńs post-hoc test for pairwise comparisons were conducted. For sample sets with normal distribution (B,C,F,H,J), ANOVA analysis with post hoc Student-Newman Keuls test were conducted. In panels B,C,F, P**<0.01 or P***<0.001 vs 0 min, P^†††^<0.01 vs 60 min. Panels B,C,E,F,H,J display means±SD as indicated.

Since previous [18–21] and current (Fig 1A-G) studies have shown that functional cytoskeleton is required for NETosis induction in the early stage of the process, and PKCα nuclear translocation requires intact cytoskeleton [16], we thought to examine the dynamic changes of actin polymerization. Previous studies reported that there is basal F-actin polymerization in the unstimulated resting cells [40, 41], and actin reorganization could be started in a “second” after cell stimulation [41, 42]. In the current study, we also found basal F-actin assembly in the unstimulated resting neutrophils (Fig 2D). Following PMA stimulation, we found a time-dependently increased percentage of cells with actin polymerization and F-actin filament formation at 15 and 30 min, reached a peak by 60 min time point, following by gradual actin depolymerization at 120 and 180 min (Fig 2D,E). In line with these observation, previous studies reported that PMA can induce actin polymerization in neutrophils and lymphocytes [37, 43]. Our results suggest that PMA-induced actin polymerization provides structural basis for nuclear translocation of PKCα [16] within the first 60 min (Fig 2D,E). While actin cytoskeleton disassembly at 120 and 180 min (Fig 2D,E) might be responsible for the plasma membrane rupture and DNA extracellular release in the corresponding time points (Fig 2A), similar to the phenomenon reported by Thiam and coauthors [22], as cortical cytoskeletal networks are important for plasma membrane integrity [10].

Interestingly, PMA stimulation increased ROCK activity at 15 and 30 min, and peaked at 60 min time point (Fig 2F), which are tightly matched with the dynamic changes of F-actin polymerization within 60 min of PMA stimulation (Fig 2D,E). Then, ROCK activity was gradually reduced, with significant decrease at 180 min as compared to that at 60 min (Fig 2F), corresponding to the gradual disassembly of actin cytoskeleton at 2-3h time points (Fig 2D,E). Therefore, the dynamic changes of ROCK activity (Fig 2F) and the corresponding actin polymerization/depolymerization (Fig 2D,E) may help to explain nuclear translocation of PKCα by functional cytoskeleton in the early stage, and plasma membrane rupture with actin depolymerization [22] in the late stage of NETosis. Therefore, actin cytoskeleton and its upstream regulator ROCK play an important role in NETosis.

Since Rho and its effector kinase ROCK regulate actin cytoskeleton organization in previous studies [23, 24], here we tested the role of ROCK in actin polymerization in neutrophils during NETosis. We found that inhibition of ROCK activation by ROCK inhibitor Y27632 resulted in impaired actin polymerization not only in primary mouse BM neutrophils (Fig 2G,H), but also in human dPMNs (Suppl Fig 1C). Additionally, genetic deficiency of ROCK1 attenuated actin polymerization in neutrophils from ROCK1 deficient mice as compared to those from WT mice (Fig 2I,J). Similarly, Gallo et al reported that ROCK1 inhibition impaired actin polymerization in cells transfected by silencing shRNA [44]. Therefore, the above results by pharmacological or shRNA silencing inhibition, or genetic deficiency, of ROCK from our and other studies [44] suggest a role of ROCK in actin polymerization. Furthermore, inhibition of ROCK by different inhibitors increased cytosolic retention of PKCα, and attenuated PKCα nuclear translocation in neutrophils (Fig 2K, Suppl Fig 1A,B), decreasing nuclear envelope disintegration (Fig 2K). Therefore, our and other studies support the role of actin cytoskeleton [16] and its upstream ROCK/MLCK/ myosin pathway (Fig 1D-F, 2K, Suppl Fig 1A,B) in nuclear translocation PKCα that is crucial to nuclear envelope rupture and extracellular release of nuclear DNA during NETosis [9, 10].

Taken together, our results demonstrated that PMA-induced reorganization of actin cytoskeleton and its upstream ROCK/MLCK/ myosin pathway (Fig 1-2) are involved in nuclear translocation of lamin kinase PKCα [16], which drives nuclear envelope rupture, leading to nuclear DNA extrusion [9, 10]. Furthermore, the reduced ROCK activity and actin depolymerization in the late stage (Fig 2D-F) may be responsible for plasma membrane rupture [22], leading to DNA extracellular release and NET formation [10]. In line with our study, several previous works also reported the involvement of actin cytoskeleton in NETosis [18–21]. The current work provides more detailed investigation and mechanistic explanation of NETosis.

### Inhibition of ROCK by pharmacological inhibition or genetic ROCK1 deficiency attenuated NET formation in neutrophils *in vitro* or *ex vivo*

To explore the role of ROCK in NETosis, we found that inhibition of ROCK with different chemical inhibitors (Y27632, HA1077, AS1892802) significantly decreased NET formation *in vitro* in dPMNs (Fig 3A) and primary human neutrophils (Fig 3B) stimulated by PMA, or in mouse neutrophils stimulated by PAF (Fig 3C), detected by different approaches. Most importantly, genetic deficiency of ROCK1 inhibited NETosis *ex vivo* in BM neutrophils from ROCK1^+/-^ mice as compared to those from their WT littermates (Fig 3D,E). Taken together, ROCK activation (Fig 2F) regulated actin polymerization (Fig 2D,E), mediating nuclear translocation of PKCα and driving nuclear envelope rupture (Fig 2A), leading to nuclear DNA extrusion. Importantly, inhibition of ROCK impaired actin polymerization (Fig 2G-J), decreased PKCα nuclear accumulation (Fig 2K), and attenuated NETosis in neutrophils *in vitro* or *ex vivo* (Fig 3A-E). Therefore, these results from *in vitro* and *ex vivo* experiments indicate that ROCK signal pathway regulated NETosis by modulation of nuclear translocation of PKCα and nuclear envelope rupture through regulation of actomyosin cytoskeletal networks (Suppl Fig 1D).

**Figure 3.**
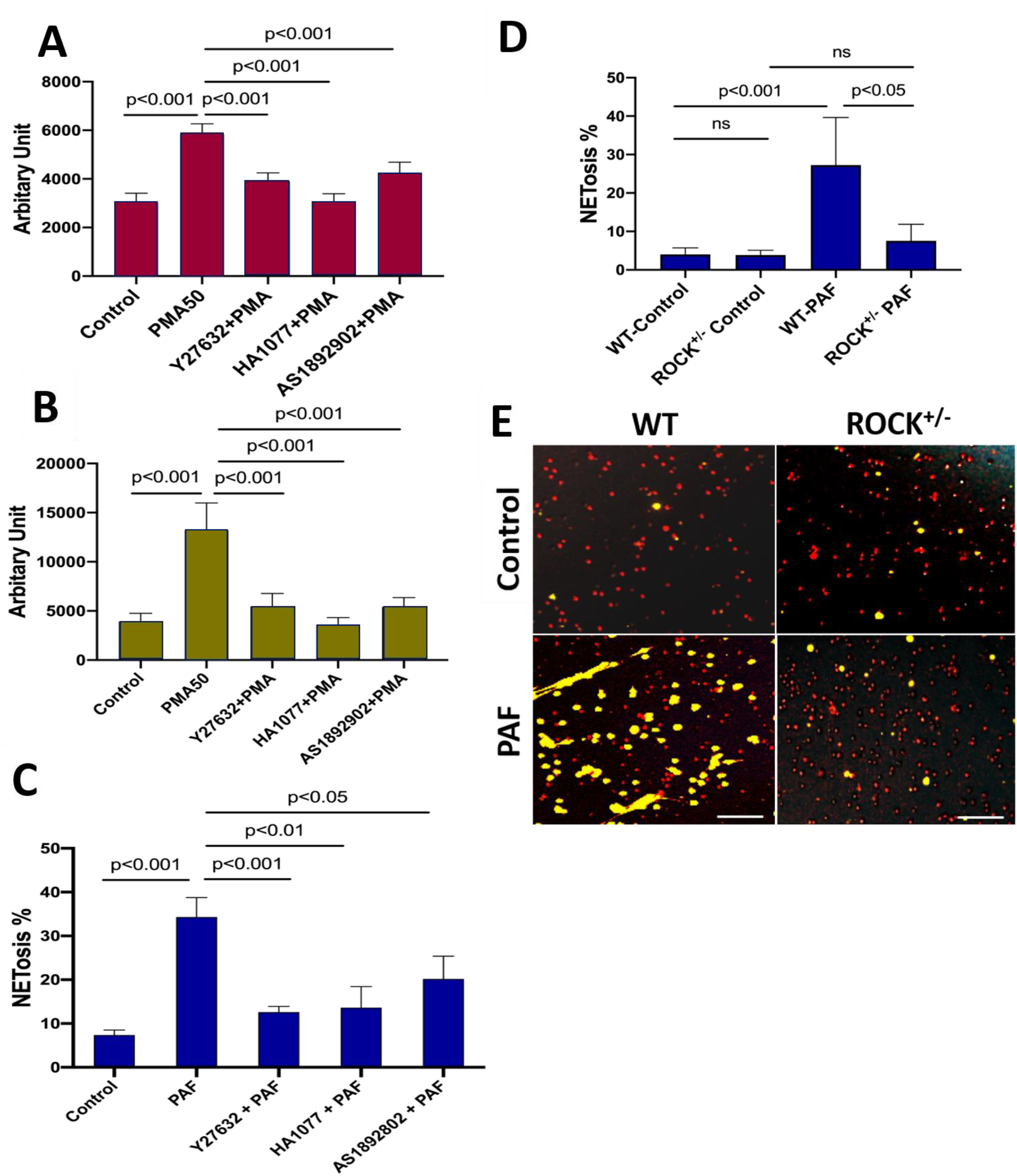
Inhibition of ROCK1 attenuated NETosis in neutrophils. Panels **A,B** show the results of NETosis of either dPMNs (**A**) or primary human neutrophils from different healthy donors (**B**), which were exposed to 50 nM PMA for 3h without or with 30min pretreatment by 50 μM ROCK inhibitors (Y27632, HA1077, AS1892802), then detection of NET formation by microplate reader. **C**-**E**, Summary analysis (**C-D**) and (**E**) representative of NET formation of mouse BM neutrophils from WT mice without or with pretreatment by different ROCK inhibitors (**C**), or BM neutrophils from WT vs ROCK1^+/-^ mice (**D-E**), and stimulated without or with 10 µM PAF (**C**,**D**,**E**) for 3h, and stained with SYTO Red and Sytox Green. Images were taken by fluorescent microscopy, and % cells with NET formation was analyzed as described in Fig 1. All results represent 5 or 6 biological replicates. Panel E represent images of the replicates (D). Scale bars, 100μm. Panels A- D display mean±SD. For non-normal distributed data (D), Kruskall-Wallis analysis and Dunńs post-hoc test were conducted. For sample sets with normal distribution (A-C), ANOVA analysis with post hoc Student-Newman Keuls test for pairwise comparisons was conducted.

### Hematopoietic-specific ROCK1 deficiency protected mice from UVB-induced skin inflammation

To study the effects of ROCK1 on neutrophil NETosis *in vivo* and its involvement in UVB-induced skin inflammation, we generated mice with hematopoietic-specific ROCK1 deficiency by transplantation of BM HSCs from ROCK1^+/-^ mice or their WT littermates to CD45.1 recipient mice, following by irradiation of these mice without or with UVB (Suppl Fig 2A). Four-weeks after the BMT procedure, the reconstitution status of donor-derived cells (CD45.2) in peripheral blood of recipient CD45.1 mice was examined by flow cytometry for each BMT mouse. Only the recipient mice with over 95% reconstitution rates (Suppl Fig 2B) were included for the following studies. To further confirm high reconstitution in mice with successful BMT, we found a majority of infiltrated neutrophils in skin lesions of UVB-irradiated CD45.1 recipient mice who received BMT of HSCs from WT donors were donor-derived CD45.2 neutrophils (Suppl Fig 2C), but not the host CD45.1 neutrophils (Suppl Fig 2D).

With these mice, we found decreased dermal skin thickness, and reduced inflammatory cell infiltration in dermal skin of UVB-irradiated BMT-ROCK1^+/-^ mice as compared to those in BMT- WT control mice (Fig 4A,C,D), however, the differences of epidermal skin thickness in BMT- ROCK1^+/-^ vs BMT-WT mice were not significant (Fig 4B). Interestingly, inflammatory cell infiltration in skin was positively correlated with dermal skin thickness in all experimental mice (Fig 4E). Furthermore, fluorescent staining of skin sections of these mice demonstrated decreased NET formation *in vivo* in skin lesions of UVB-irradiated BMT-ROCK1^+/-^ mice as compared to those of BMT-WT control mice (Fig 4F,G). Importantly, distribution area of neutrophil NETs in inflamed skin was positively correlated with dermal skin thickness (Fig 4H), indicating the involvement of neutrophil NETs in skin inflammation.

**Figure 4.**
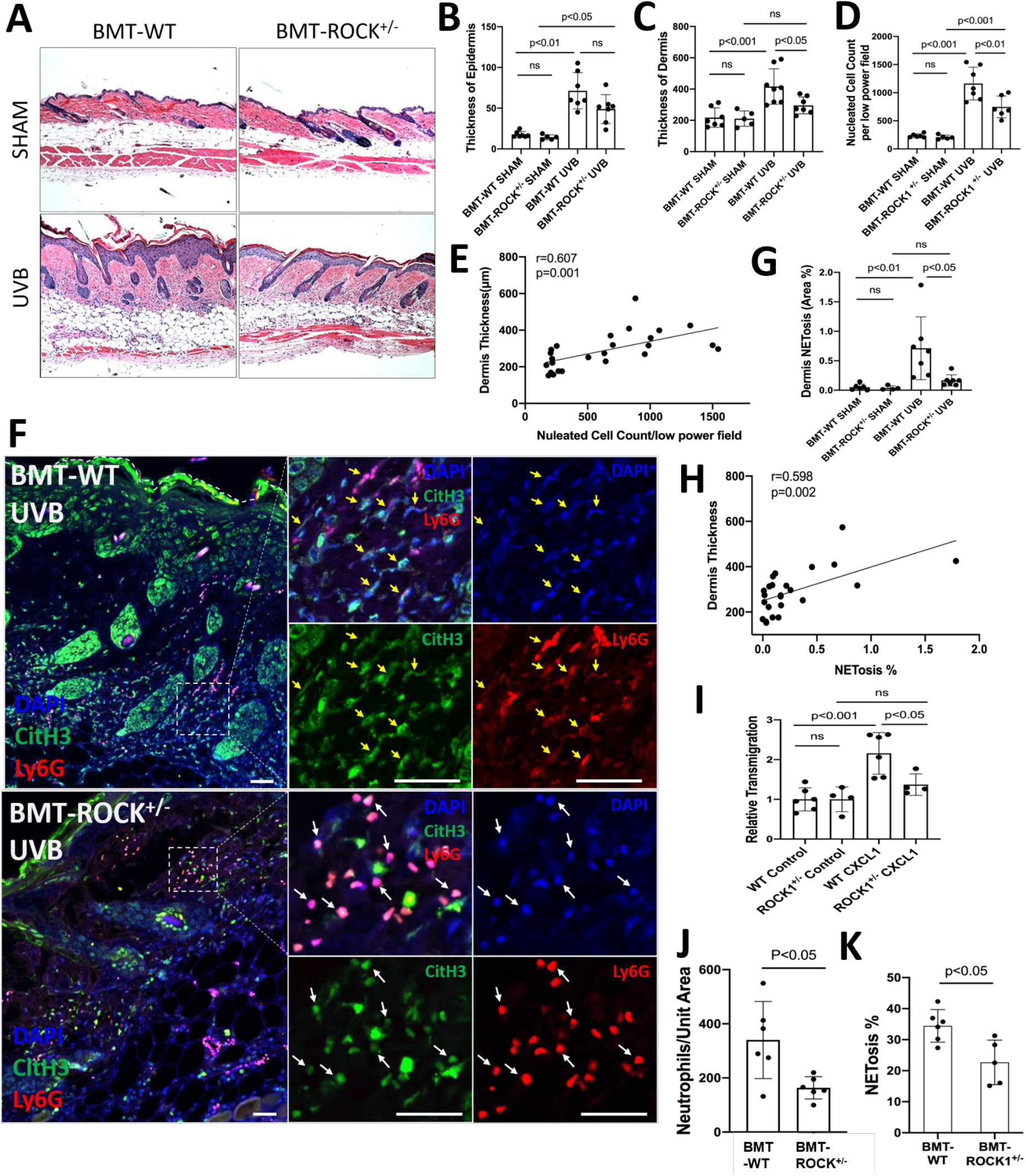
Hematopoietic-specific ROCK1 deficiency protected mice from UVB-induced skin inflammation in BMT-ROCK1^+/-^ mice. **A-E**. Representative H&E staining images (A) and summary analysis of the thickness of epidermis and dermis (B,C), infiltration of inflammatory (nucleated) cells in the dermis and subcutis (D) and their correlation with dermis thickness (E) based on H&E staining of skin sections of BMT-WT vs BMT-ROCK1^+/-^ mice that were irradiated without (sham) or with UVB. **F-H.** Representative images (F) and summary analysis of NETosis (G), and its correlation with skin thickness (H) in skin of BMT-WT vs BMT-ROCK1^+/-^ mice that were irradiated without (sham) or with UVB. **I**. Relative migration analysis of BM neutrophils of WT vs ROCK1^+/-^ mice was conducted with Boyden chamber by stimulation without or with CXCL1. **J**. Neutrophil infiltration per unit area in skin of UVB-irradiated BMT-WT vs BMT- ROCK1^+/-^ mice. **K**. Percentage of NETotic cells among neutrophils in skin of UVB-irradiated BMT-WT vs BMT-ROCK1^+/-^ mice. For panels F-H,J,K, skin sections were stained by antibodies against **Ly6G** (PE) neutrophil marker, citrullinated histone H3 (**citH3**, FITC), and **DAPI** for DNA. In panels A,F, Scale bars, 100 μm. All results represent 4-8 biological replicates as indicated in individual panels. For non-normal distributed data sets (B), Kruskall-Wallis analysis and Dunńs post-hoc pairwise comparisons were conducted. For sample sets with normal distribution (C,D,G,I,J,K), multiple comparisons were conducted by ANOVA analysis with Student-Newman Keuls post hoc pairwise comparisons, while student’s t-test were conducted for two groups comparisons (J,K).

To understand the role of ROCK1 deficiency in skin inflammation with neutrophil NETosis, we found mildly decreased chemotactic migration ability *in vitro* in ROCK1 deficient neutrophils as compared to that in WT neutrophils (Fig 4I). A previous study by our co-author reported that ROCK1 deletion reduced CXCL1/CXCR2-mediated migration in macrophages derived from bone marrow of ROCK1 deficient mice [45]. Furthermore, no differences in CXCR2 expression in BM neutrophils from WT vs ROCK1^+/-^ mice were observed (Suppl Fig 2E,F). Generally, cell migration is affected by cellular factors (including cytoskeleton) and environmental factors (including chemotactic signals) [46]. The above results indicate an effect of destabilized actin cytoskeleton on impaired chemotactic migration ability in ROCK1 deficient neutrophils. These results may explain in part the reduced infiltration of inflammatory cells (Fig 4D), particularly neutrophils (Fig 4J), in inflamed skin of UVB-irradiated BMT-ROCK1^+/-^ mice as compared to those in BMT-WT controls. However, the mildly reduced chemotactic migration ability of heterogeneous ROCK1 deficient neutrophils in vitro seems insufficient to explain the ameliorated skin inflammation in UVB-irradiated BMT-ROCK1^+/-^ mice as compared to those in their BMT-WT controls (Fig 4A-D,J).

Recent studies demonstrated that NETs have chemotactic function to promote cancer cell metastasis [47] or enhance monocyte infiltration by induction of MCP1 expression in endothelial cells [48]. Here, we found decreased NETosis *in vitro* in human or mouse neutrophils with ROCK inhibitors (Fig 3A-C), and *ex vivo* in neutrophils from ROCK1^+/-^ mice (Fig 3D,E). Most importantly, we found a significantly reduced percentage of NET formation *in vivo* among neutrophils infiltrated to the inflamed skin of UVB-irradiated BMT-ROCK1^+/-^ mice as compared to those in BMT-WT mice (Fig 4K). Therefore, the attenuated skin inflammation in UVB-irradiated BMT-ROCK1^+/-^ mice may, at least in part, be attributed to the reduced NET formation by ROCK1 deficiency (Fig 4A,F,G). Our published work has shown that neutrophil NETs can display IL-17A and TNFα, cytokines with chemotactic properties, in the skin of UVB-irradiated mice [9]. Next, we sought to look the effects of ROCK1 deficiency on display of NET-associated cytokines in skin of UVB-irradiated BMT-ROCK1^+/-^ mice.

### Hematopoietic-specific ROCK1 deficiency decreased the exhibition of pro-inflammatory cytokines by neutrophil NETs in BMT-ROCK1^+/-^ mice

We found that UVB exposure increased formation of NETs with exhibition of NET-associated IL-17A, TNFα, IFNγ, or IFNα in inflamed skin of BMT-WT mice (Fig 5A-I). Importantly, hematopoietic-specific ROCK1 deficiency attenuated neutrophil NETosis *in vivo* and its display of NET-associated cytokines in UVB-irradiated skin of BMT-ROCK1^+/-^ mice (Fig 5A- I). Many cytokines have chemotactic properties for recruitment of inflammatory cells [49–51]. Here, we found NET-associated IL-17A or TNFα in inflamed skin in UVB-irradiated BMT-WT mice (Fig 5A-I) similar to what we found in UVB-irradiated mice in our recent publication [9]. Previous studies have reported that IL-17A [49], together with TNFα [50], can synergistically induce leukocyte infiltration to sites of inflammation [52]. Furthermore, NET-associated IFNγ together with IL-17A also promoted leukocyte recruitment [51]. Thus, extracellular display of NET-associated IL-17A, TNFα, and IFNγ in skin contributes to further recruitment of leukocytes and their accumulation in certain areas of the dermis, as well as propagation of inflammatory responses in inflamed skin of UVB-irradiated BMT-WT mice (Fig 4F, 5C). Importantly, there were fewer and diffusely distributed non-netting leukocytes in skin of UVB-irradiated BMT- ROCK1^+/-^ mice (Fig 4F, 5C). After UVB irradiation, the neutrophil NETs and their associated cytokines with relatively high concentrations in the skin likely contributed to further recruitment of leukocytes to the skin of BMT-WT mice. In contrast, the non-netting neutrophils may secrete their cytokines to the extracellular space that always quickly diffused to the bloodstream, where they are diluted by the circulation in UVB-irradiated BMT-ROCK1^+/-^ mice. Therefore, this may partly explain the decreased leukocyte infiltration in skin of UVB-irradiated BMT-ROCK1^+/-^ mice (Fig 4F, 5C). In addition to above cytokines, we also detected NET-associated IFNα, an important cytokine that is crucial to pathogenesis of SLE [53, 54] and important to UVB-induced skin inflammation [33]. More importantly, NET-associated IL-17A, TNFα, IFNγ, and IFNα in skin correlated with dermal skin thickness in all experimental mice (Suppl Fig 3).

**Figure 5.**
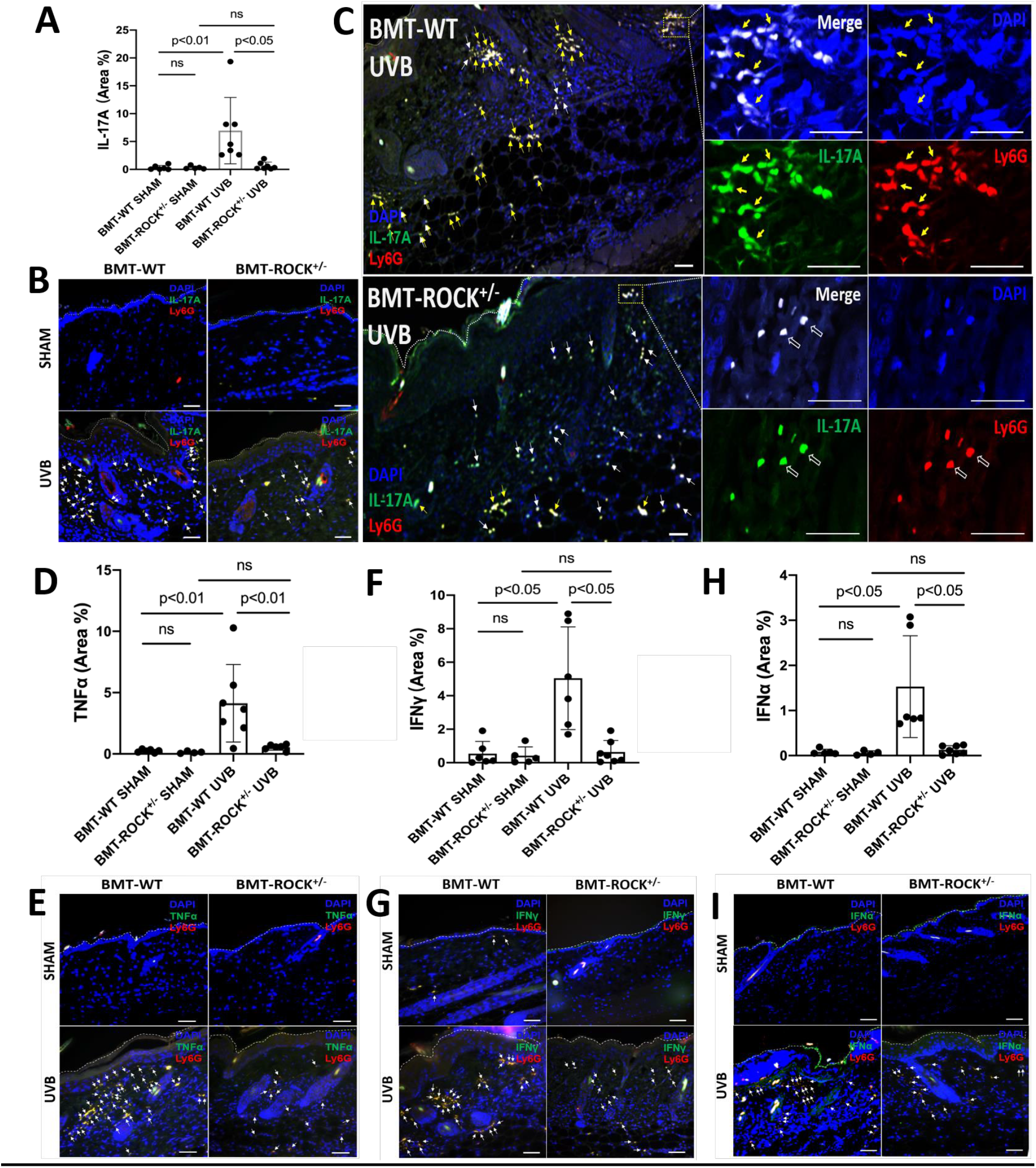
Hematopoietic-specific ROCK1 deficiency attenuated skin inflammation with decreased exhibition of NET-associated pro-inflammatory cytokines in UVB-irradiated BMT-ROCK1^+/-^ mice. **A-I**. Summary analyses (**A**,**D**,**F**,**H**) and representative images (**B**,**C**,**E**,**G**,**I**) of skin sections of BMT-WT vs BMT-ROCK1^+/-^ mice that were irradiated without (sham) or with UVB, and stained by PE-conjugated rat anti-mouse **Ly6G** Ab, and FITC labeled rat anti-mouse **IL-17A**, **TNFα**, **IFNγ**, or **IFNα**. DNA was stained by **DAPI** for panels **A**-**I**. Yellow arrows indicate NETotic neutrophils, and white arrows show non-netting neutrophils in panels B,C,E,G,I. Scale bars, 100μm. All results represent 4-7 biological replicates as indicated in individual panels (A,D,F,H). For non-normal distributed data sets (A,F), Kruskall-Wallis analysis and Dunńs post- hoc pairwise comparisons were conducted. For sample sets with normal distribution (D,H), multiple comparisons were conducted by ANOVA analysis with Student-Newman Keuls post hoc pairwise comparisons.

### ROCK1 genetic deficiency inhibited formation of NETs *ex vivo* and their display of pro-inflammatory cytokines in neutrophils from ROCK1 deficient mice

Previous publications have reported that neutrophils can express and produce various cytokines, including IL-17A [51], TNFα [55], IFNγ [56], or IFNα [57]. Studies from our and other groups have reported that neutrophil-expressed cytokines can be released with NETs [9, 57]. Recently, Agak et al demonstrated that extracellular traps released by TH17 cells can also be associated with their expressed IL-17 [58]. In the current study, we found the display of NET-associated cytokines in UVB-irradiated skin (Fig 5). To determine if these NET-associated cytokines were derived from their parental neutrophils, we conducted *ex vivo* experiments by treatment of mouse neutrophils with PAF, a physiologic NETosis inducer that can be released from keratinocytes after UVB irradiation [9, 59]. We found that PAF stimulation can induce expression of IL-17A, TNFα, IFNγ, and IFNα (with PAF+NETs) in mouse BM neutrophils, detected by flow cytometry analyses (Fig 6A,D,G,J, Suppl Fig 4). Interestingly, PAF-induced IL-17A, TNFα, IFNγ, and IFNα in mouse neutrophils can be released from their parental neutrophils to the extracellular space during NET formation (Fig 6B,C,E,F,H,I,K,L).

**Figure 6.**
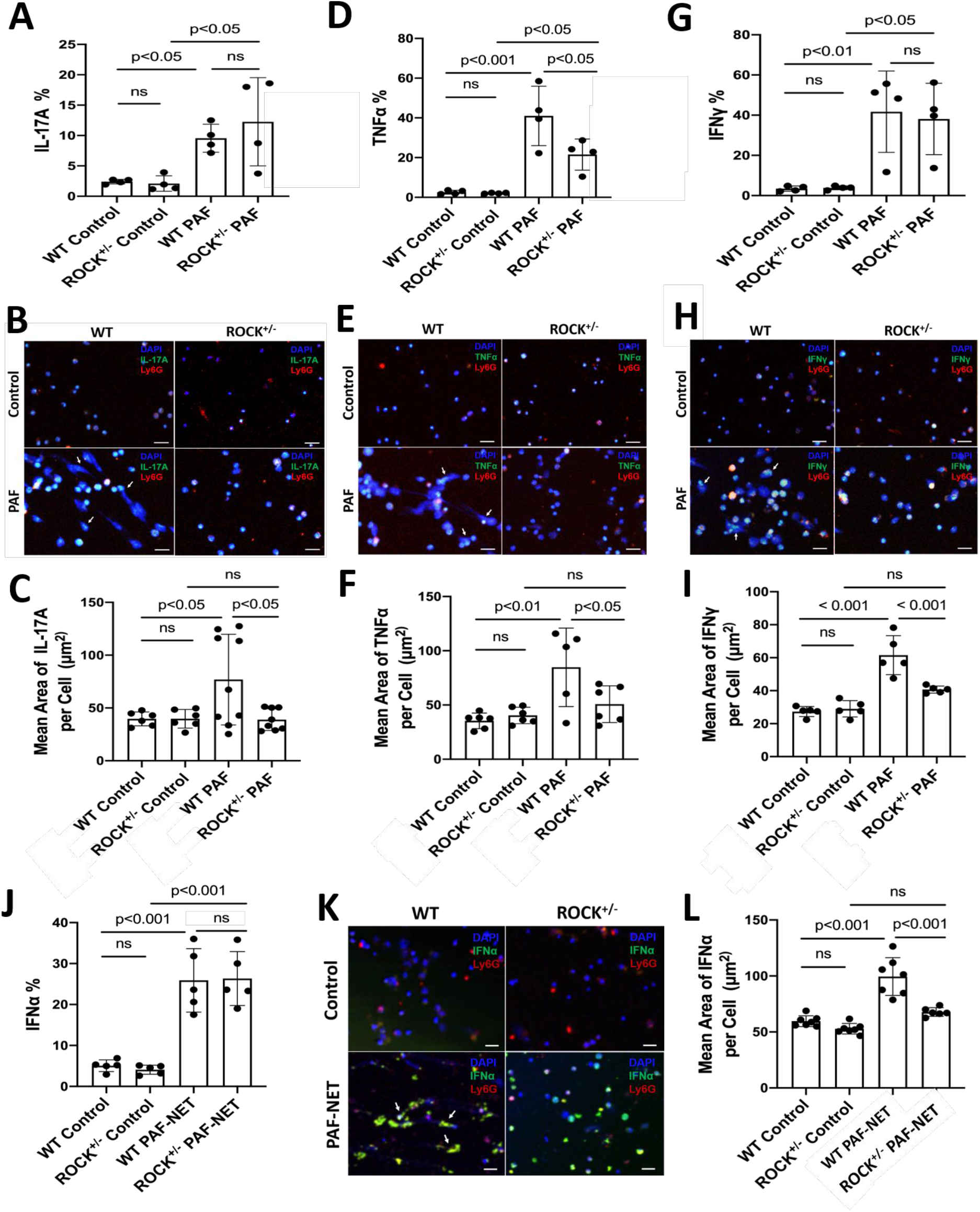
Genetic deficiency of ROCK1 decreased formation of NETs and attenuated their display of pro-inflammatory cytokines in neutrophils from ROCK1 deficient mice. **A**-**L**. Summary flow cytometry analyses (**A**,**D**,**G,J**), representative fluorescent images (**B**,**E**,**H,K**) and their summary analyses (**C**,**F**,**I**,**L**) of BM neutrophils from ROCK1^+/-^ mice or their WT littermates were treated without or with 10 µM PAF (for IFNα induction in **J**-**L**, treatment with PAF and isolated NETs from cultured neutrophils of MRL/lpr mice) for 20h, followed by fixation, permeabilization, and staining of neutrophils by PE-conjugated anti-mouse **Ly6G**, FITC labeled anti-mouse **IL-17A**, **TNFα**, **IFNγ**, **or IFNα**, then following by flow cytometry or confocal microscopy analyses. DNA was stained by **DAPI.** Scale bars, 50μm. Arrows indicate NETs with extracellular display of different cytokines. All results represent 4-9 biological replicates as indicated in individual panels. All sample sets with normal distribution, the multiple comparisons were conducted by ANOVA analysis with Student-Newman Keuls post hoc tests.

ROCK1 deficiency attenuated PAF-induced TNFα (Fig 6D, Suppl Fig 4B), but did not affect expression of IL-17A, IFNγ, and IFNα (Fig 6A,G,J, Suppl Fig 4) in BM neutrophils from ROCK1^+/-^ mice as compared to those from WT mice. Importantly, ROCK1 deficiency attenuated NET formation with decreased area of extracellular display of NET-associated cytokines by enclosing them within non-netting neutrophils from ROCK1^+/-^ mice as compared to those from WT mice (Fig 6B,C,E,F,H,I,K,L). After UVB irradiation, PAF from keratinocytes can trigger neutrophils to form NETs that exhibit proinflammatory cytokines, which further recruited leukocytes, therefore resulting in increased inflammatory cell accumulation in skin, particularly in the epidermal-dermal junction area of inflamed skin (Fig 4A,F,5C). Although ROCK1 deficiency did not affect cellular expression of some cytokines (Fig 6A,D,G,J, Suppl Fig 4), inhibition of NET formation by ROCK1 deficiency did decrease their extracellular display with NETs by increasing their intracellular retention, thus resulting in reduced leukocyte recruitment in skin with ameliorated skin inflammation in UVB-irradiated BMT-ROCK1^+/-^ mice.

## DISCUSSION

Photosensitive skin reaction is a complex inflammatory response to UVB-induced skin tissue damage [2, 4, 60, 61]. UVB overexposure causes epidermal keratinocyte apoptosis and DNA damage [61, 62], and accumulation of dead cell extracellular DNA may initiate autoimmune responses for lupus development [4]. In addition, other types of cell death have also been shown to contribute to extracellular DNA accumulation, and lupus pathogenesis [63]. Histopathologic analysis of skin lupus indicates the potential interaction between immune cells and epidermal keratinocytes [61, 64]. UVB irradiation of keratinocytes releases PAF, which contributes to the sustained inflammatory responses [35], inducing neutrophil recruitment and NETosis in inflamed skin [9]. Studies from our and other groups [9, 59] have shown that PAF can serve as a natural stimulus for induction of NETosis. On the other hand, direct exposure of neutrophils to UV can also induce NET formation *in vitro* [9, 65]. Therefore, NET formation *in vivo* in UVB-irradiated skin may be attributed to indirect effects of UVB through PAF [35] and direct effects of UVB irradiation [9, 65] on neutrophils infiltrated to inflamed skin.

Over the past decade, various signaling pathways have been reported to be involved in NETosis, i.e., NADPH oxidase (NOX)-dependent vs -independent, reactive oxygen species (ROS)-dependent vs -independent [66–68]. Given that nuclear chromatin forms the backbone of NETs, the nucleus is the root of nuclear DNA extracellular traps [10, 69–73]. Thus, decondensation of nuclear chromatin is required for its externalization, while rupture of the nuclear envelope and plasma membrane are the necessary steps for removal of the physical barriers of nuclear membranes for nuclear DNA discharge and extracellular NET formation [10]. Therefore, these key cellular morphological changes in nuclear chromatin, nuclear envelope, and plasma membrane are required for extracellular DNA NET formation [9, 10, 14, 72, 73] no matter if NETosis is NOX- or ROS-dependent or –independent.

We recently reported that NETotic nuclear envelope rupture is driven by PKCα-mediated nuclear lamin B disassembly [9]. In the current study, we found that nuclear translocation of cytosolic PKCα [9, 15] was regulated by actin cytoskeleton [16], actomyosin cytoskeletal networks and their upstream ROCK in the early stage of NETosis induction. In line, Hippenstiel demonstrated that inhibition of ROCK blocks nuclear translocation and activation of PKC in endothelial cells [17], and Matoba reported the role of ROCK in nuclear translocation of other cytosolic molecules through regulation of actin cytoskeleton organization [74]. Other studies also report the involvement of actin cytoskeleton filament [38], or myosin II [39], or actomyosin-dependent [25] in nucleocytoplasmic shuttling. Therefore, the current study provides a cellular mechanistic explanation for the role of actin cytoskeleton in early stage of NETosis induction [18, 21], in which nuclear translocation of NETotic lamin kinase PKCα is mediated by actin cytoskeleton [16], and its upstream ROCK signaling pathway [17] during NETosis. In addition to their role in nuclear-cytoplasmic shuttling [25, 26], ROCK and its-regulated actomyosin cytoskeletal networks may also enable nuclear deformation and contribute to nuclear envelope rupture by forces generated by the cytoskeletal networks [75]. However, it is still unclear if actomyosin cytoskeletal networks also contribute to NETotic nuclear envelope rupture by tearing the disassembled nuclear lamina with their “forces”.

Plasma membrane is the final physical barrier for extracellular release of nuclear chromatin and NET formation. The cell cortex is a layer of cytoskeletal networks underneath the plasma membrane, consisting of F-actin filaments, myosin motors, and actin-binding proteins [76]. The contractile forces generated by cortical cytoskeletal networks are important for maintenance of plasma membrane integrity [76]. In contrast to ROCK activation and actin cytoskeleton polymerization in the early stage of NETosis induction, their dynamic changes after the peak time point (60 min) turned to gradually decrease in ROCK activity with actin cytoskeleton disassembly during the late stage of NETosis induction. The late stage actin cytoskeleton disassembly may contribute to the plasma membrane rupture for NET release as reported in a recent study [22].

UVB-induced skin inflammation manifested as increased skin thickness, enhanced inflammatory cell infiltration, and elevated proinflammatory cytokine expression. ROCK and its regulated actin cytoskeletal networks serve as cellular factors to modulate cell migration [46]. Wang and colleagues demonstrated that macrophages with ROCK1 deletion from ROCK1^-/-^ mice showed significantly impaired chemotactic migration ability [45]. In the current study, we found the mildly decreased transmigration *ex vivo* in transwell migration experiments in neutrophils with heterogeneous ROCK1 deficiency, and the significantly reduced leukocyte infiltration *in vivo* with diffusely distributed non-netting neutrophils in skin of UVB-irradiated BMT-ROCK1^+/-^ mice. These results suggest that the reduced ROCK1 expression in leukocytes of ROCK1^+/-^ mice [28] can still partly regulate actomyosin cytoskeletal networks that drive cell movement. In the meantime, the reduced ROCK1 expression can inhibit NETosis in neutrophils through interfering with cytosolic PKCα nuclear translocation that induces nuclear envelope rupture [9].

We have recently demonstrated the involvement of NETs and NET-associated cytokines in UVB-induced skin inflammation [9], and others also demonstrated the association of IL-17 with neutrophil NETs [77] or extracellular traps released by TH17 cells [58]. In the current study, we found PAF-induced expression of IL-17A, TNFα, IFNγ, and IFNα in neutrophils, and these cytokines can be released with NETs to the extracellular space during NETosis. Furthermore, ROCK1 deficiency not only inhibited TNFα expression in neutrophils, but also attenuated NETosis and TNFα release with NETs ex vivo, thus contributing to decreased skin inflammation in UVB-irradiated BMT-ROCK1^+/-^ mice. Several other studies have shown that inhibition of ROCK can attenuate TNFα expression in different cells in various scenario [78, 79]. However, we did not see significant effects of ROCK1 deficiency on neutrophil expression of IL-17A, IFNγ and IFNα cytokines in the current study. Several papers have reported that the ROCK2 isoform, but not ROCK1, regulates IL-17 and IFN-γ secretion from T cells [80, 81], as IL-17 production can be inhibited by either ROCK2 genetic deficiency [80], or ROCK2 siRNA silencing or ROCK2 specific inhibitor KD025 [81]. In addition, KD025 can attenuate IFN-γ production, while ROCK1 siRNA silencing does not affect IFN-γ production [81]. These studies help explain our findings of unchanged IL-17A and IFNγ in ROCK1 deficient neutrophils as compared to the WT cells. However, there is no published study regarding the effects of ROCK on IFNα expression. Although no changes were observed in neutrophil expression of several cytokines, ROCK1 deficiency can inhibit NET formation and decrease extracellular display of NET-associated IL- 17A, IFNγ and IFNα ex vivo, and in vivo in skin of UVB-irradiated BMT-ROCK1^+/-^ mice. Taken together, decreased extracellular exhibition of NET-associated IL-17A, TNFα, IFNγ, and IFNα may be responsible for the attenuated leukocyte recruitment and skin inflammation in UVB-irradiated BMT-ROCK1^+/-^ mice.

Using mice with genetic deficiency, Li et al reported that both IL-17A and IFN-γ can be produced by neutrophils and can positively regulate neutrophil infiltration [51]. Importantly, we recently demonstrated that blockage of TNFα with anti-TNFα significantly inhibited UVB-induced recruitment of inflammatory cells [82]. In contrast to the freely secreted cytokines that can be diffused to circulation with large dilution by circulating blood, the NET-associated cytokines may be retained in the local skin at a relatively high concentration for longer period of time, thus sustaining and propagating inflammatory responses in UVB-irradiated skin. All of the above support the role of NET-associated IL-17A, TNFα, or IFNγ cytokines in inflammatory cell recruitment to skin of UVB irradiated mice, while inhibition of NET formation with decreased extracellular display of NET-associated cytokines may at least partly explain the attenuated infiltration of leukocytes in inflamed skin of BMT-ROCK1^+/-^ mice. In line with the current study, recent publications demonstrated that NETs can act as a chemotactic factor with NET-associated components, i.e. human LL-37 [83] or mouse CRAMP [84], to promote cell migration, including cancer cell metastasis [47]. On the other hand, NETs may enhance monocyte infiltration by induction of MCP1 expression in endothelial cells [48].

In addition to the above discussed chemotactic effects of NET-associated cytokines, NET-associated IFNγ and IFNα not only contribute to UVB-induced skin inflammation [61, 85], but are also involved in lupus pathogenesis [86–88]. Although plasmacytoid dendritic cells are professional cells for IFNα production, recent works indicate that neutrophils may also express IFNα in the presence of chromatin [57]. In the current study, we found NETotic neutrophils express IFNα *in vivo* in skin of UVB-irradiated BMT-WT mice, probably triggered by DNA released from NETotic neutrophils nearby in inflamed skin, while ROCK1 deficiency may decrease IFNα expression through inhibition of DNA NET exposure in skin of BMT-ROCK1^+/-^ mice. This is also confirmed by our *ex vivo* experiments in which IFNα expression in neutrophils can be produced by co-stimulation with PAF and isolated NETs. Since IFNα is a major driver of lupus pathogenesis, our results suggest that NETs and their associated IFNα may contribute to photosensitivity in lupus. Isgro et al reported enhanced ROCK activation in SLE patients with increased Th17 cell differentiation, and inhibition of ROCK attenuated IL-17 production in SLE patients [89]. Thus, ROCK might be a potential target for regulation of T cell dysfunction in SLE [90]. Interestingly, the current study demonstrated that ROCK might be a potential target for treatment of UVB-induced skin inflammation due to its role in regulation of NET formation.

Taken together, ROCK regulated NET formation in neutrophils by modulation of nuclear translocation of PKCα through actomyosin cytoskeletal networks. ROCK1 deficiency ameliorates UVB-induced skin inflammation by attenuation of NET formation and its display of NET-associated cytokines. The current study demonstrates the contribution of NETosis to UVB-induced skin inflammation and photosensitivity in lupus, thus providing insights into novel therapeutics in NETosis-related human diseases, including lupus.

## AUTHOR CONTRIBUTIONS

Liu conceived and designed the study. Li, Lyu, Liu performed the laboratory experiments and acquired the experimental data. For the microscopic image analysis, Liu, Lyu, Li took the images, Lyu and Li conducted further quantification and statistical analysis. Liu, Lyu, and Werth analyzed and interpreted the data and finalized the paper. Li, Lyu, Liao, Werth and Liu were involved in preparation and revision of the manuscript. All authors read and approved the final manuscript.

## ACKNOWLEDGEMENTS

The authors would like to acknowledge Penn Skin Biology and Diseases Resource-based Center for their skin histology service. The authors would thank Debra A. Pawlowski (animal facility, Philadelphia VA Medical Center) for her kindly help with our animal experiments. This work was supported by Lupus Research Alliance (416805) and NIH R21AI144838 (to MLL), the Veterans Affairs Merit Review Award (to VPW), and NIH HL052233 and HL136962 (to JL).

## CONFLICT OF INTEREST

The authors have no conflict of interests to declare.

## Supplemental Materials

### Supplemental Figures

**Supplemental Fig 1.**
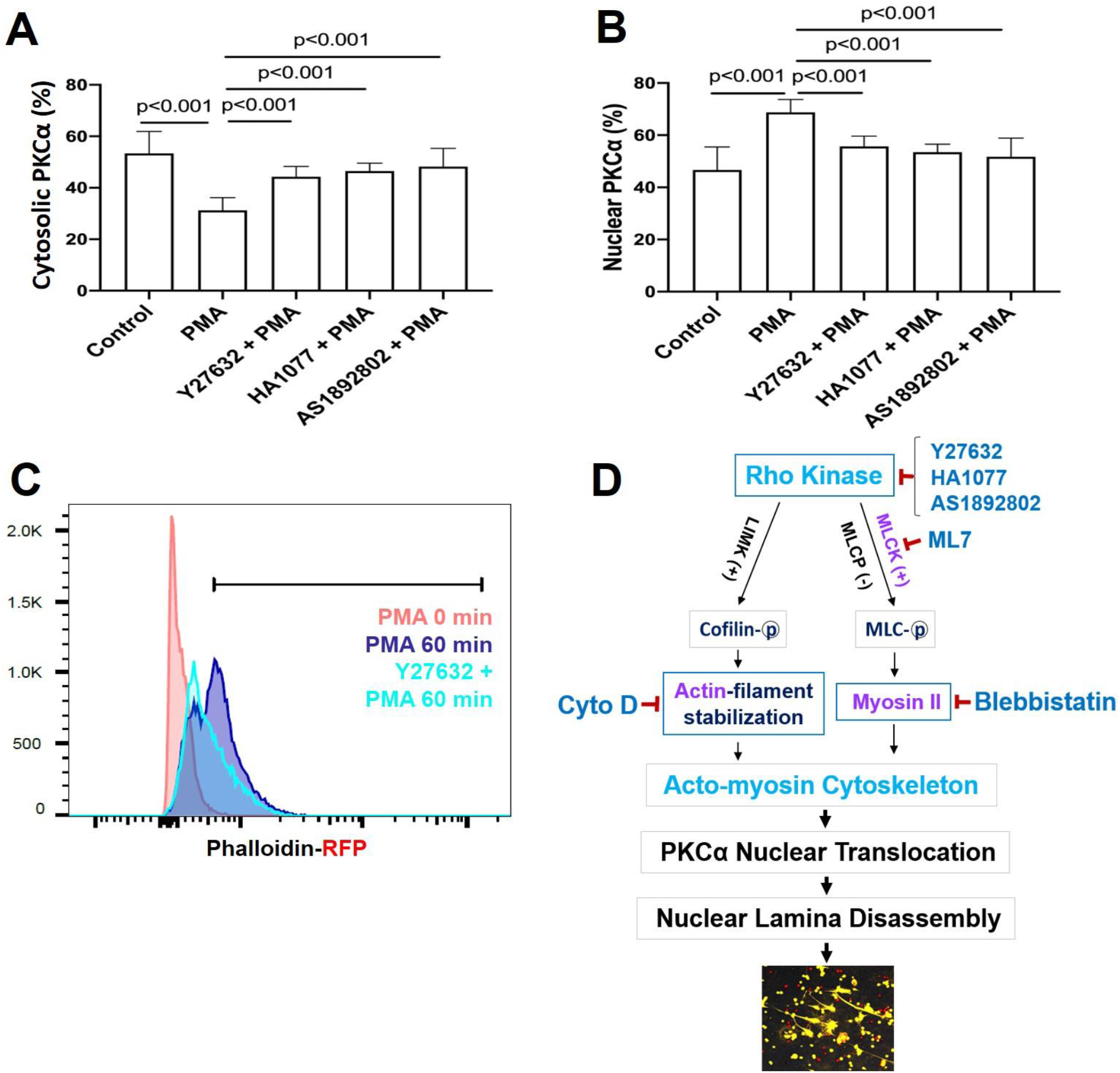
**A,B.** Summary analyses of time course study of PKCα distribution in cytosolic (A) vs nucleus (B) in dPMNs treated by 50 nM PMA for 60 min, then stained for PKCα (primary anti-PKCα, and PE-labeled secondary antibody) and lamin B (primary anti-lamin B, and FITC-labeled secondary antibody), following by confocal fluorescent microscopy, and Image J analyses. **C.** Representative analysis of F-actin polymerization detected by phalloidin-RFP with flow cytometry in dPMNs stimulated by 50 nM PMA for 60 min without or with pretreatment by 50 μM Y27632, ROCK inhibitors for 30 min. **D.** Schematic illustration of the involvement of actin cytoskeleton, actomyosin cytoskeletal networks, their upstream signaling pathways, and ROCK, as well as their corresponding inhibitors in NETosis. Panels A,B were summary analyses of at least 10 cells of each condition from three independent experiments. ANOVA analysis with post hoc Student-Newman Keuls test were conducted for these sample sets with normal distribution.

**Supplemental Fig 2.**
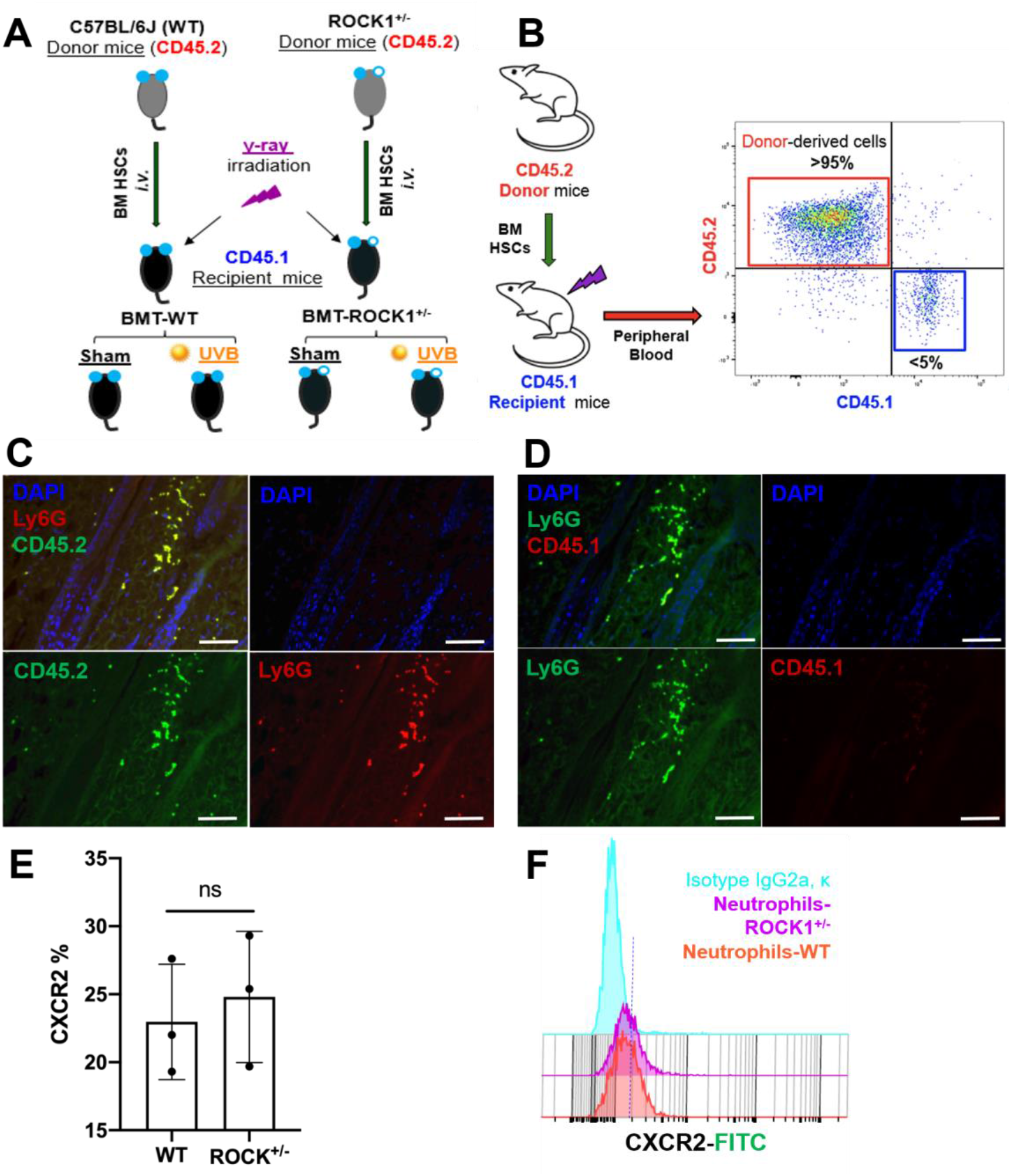
**A.** Experimental design of the BMT-WT and BMT-ROCK1^+/-^ mice that were irradiated without or with UVB. **B.** Representative dot plots of flow cytometry analysis of all BMT experimental mice for detection of reconstitution rate of donor-derived cells (CD45.2) in peripheral blood of recipient mice (CD45.1). **C,D.** Representative images of immunohistochemistry staining of skin lesions from UVB-irradiated CD45.1 recipient mice with BMT of HSCs from WT donors, stained by antibodies against Ly6G (PE) and CD45.2 (FITC) (C), or antibodies against CD45.1 (PE) and LY6G (FITC) (D), and DAPI for DNA. **E,F.** Summary analyses and representative histograms of flow cytometry analyses of CXCR2 expression in BM neutrophils from WT vs ROCK1+/- mice, stained by fluorescent conjugated antibodies against mouse Ly6G (PE), CXCR2 (FITC) and its isotype control IgG2a,κ (FITC). Un-paired student’s t-test was conducted (E), and statistical significance was considered at a level of P-value < 0.05.

**Supplemental Fig 3.**
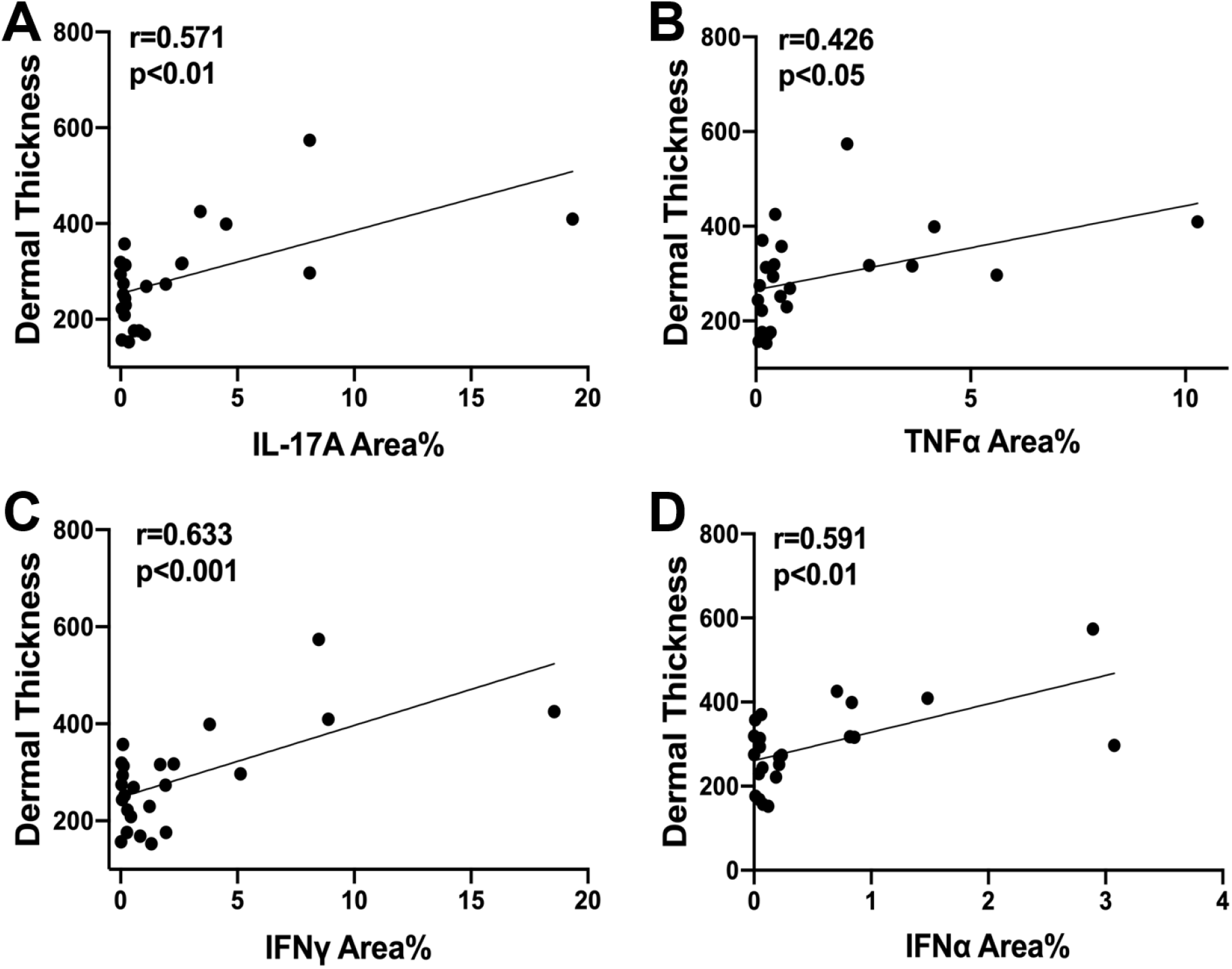
**A-D.** Correlation between NET-associated IL-17A (Area %), TNFα, IFNγ, or IFNα and dermal thickness in all groups of BMT experimental mice. Pearson correlation analysis was conducted to analyze correlation between NET-associated cytokines and dermal skin thickness.

**Supplemental Fig 4.**
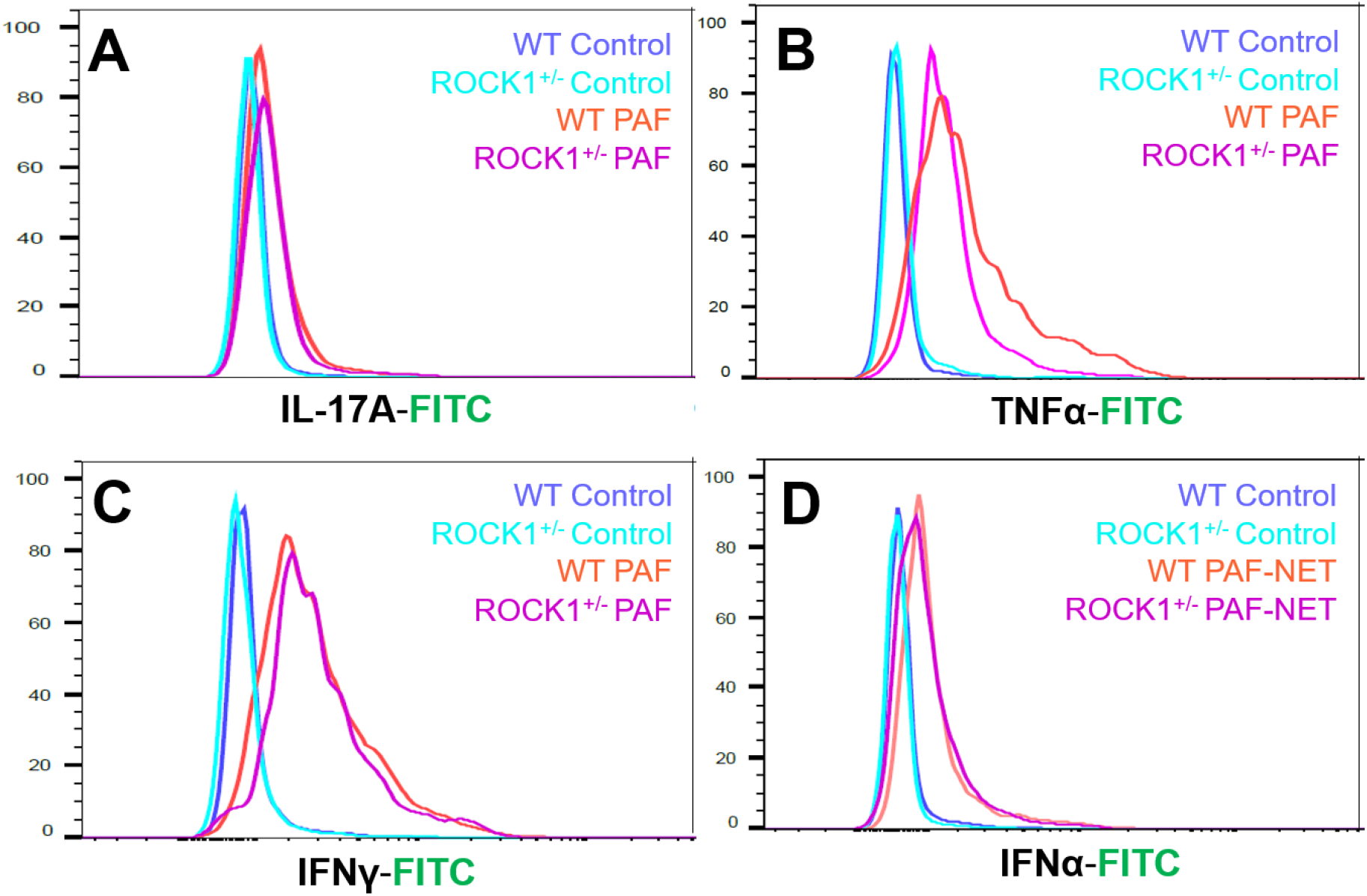
**A-D.** Representative histograms of flow cytometry analyses of BM neutrophils, from ROCK1+/- mice or their WT littermates, which were treated without or with 10 µM PAF (for IFNα induction in panel D, treatment with PAF and NETs isolated from cultured neutrophils of MRL/lpr mice) for 20h, then fixation and staining with PE-conjugated anti-mouse Ly6G, and FITC labeled anti-mouse IL-17A, TNFα, IFNγ, or IFNα, following by flow cytometry analyses (Summary data, see Fig 6A,D,G,J).

## Notes

### Competing Interest Statement

The authors have declared no competing interest.

### Summary of Updates

Li et al. Rho kinase and NETosis

